# A SARS-CoV-2 nucleocapsid protein TR-FRET assay amenable to high-throughput screening

**DOI:** 10.1101/2021.07.03.450938

**Authors:** Kirill Gorshkov, Desarey Morales Vasquez, Kevin Chiem, Chengjin Ye, Bruce Nguyen Tran, Juan Carlos de la Torre, Thomas Moran, Catherine Z. Chen, Luis Martinez-Sobrido, Wei Zheng

## Abstract

Drug development for specific antiviral agents against coronavirus disease 2019 (COVID-19) is still an unmet medical need as the pandemic continues to spread globally. Although huge efforts for drug repurposing and compound screens have put forth, only few compounds remain in late stage clinical trials. New approaches and assays are needed to accelerate COVID-19 drug discovery and development. Here we report a time-resolved fluorescence resonance energy transfer-based assay that detects the severe acute respiratory syndrome coronavirus 2 (SARS-CoV‑2) nucleocapsid protein (NP) produced in infected cells. It uses two specific anti-NP monoclonal antibodies (MAbs) conjugated to donor and acceptor fluorophores that produces a robust ratiometric signal for high throughput screening of large compound collections. Using this assay, we measured a half maximal inhibitory concentration (IC_50_) for Remdesivir of 9.3 μM against infection with SARS-CoV-2 USA/WA1/2020 (WA-1). The assay also detected SARS-CoV-2 South African (Beta, β), Brazilian/Japanese variant P.1 (Gamma, γ), and Californian (Epsilon, ε), variants of concern or interest (VoC). Therefore, this homogeneous SARS-CoV-2 NP detection assay can be used for accelerating lead compound discovery for drug development and for evaluating drug efficacy against emerging SARS-CoV-2 VoC.

## Introduction

Severe acute respiratory syndrome coronavirus 2 (SARS-CoV-2) emerged in late 2019 and has been responsible for the deadly global pandemic of coronavirus disease 2019 (COVID-19), rivaled only by the “Spanish Flu” pandemic of 1918-1919 (40-50 million deaths) and the ongoing HIV/AIDS pandemic from 1981-present (25-30 million deaths) (Visualizing the History of Pandemics, https://www.visualcapitalist.com/history-of-pandemics-deadliest/). As of June 2021, there have been over 168 million infection cases and 3.5 million deaths due to COVID-19 and these tolls are increasing daily ^1^. Thought to have been transmitted from animal to human, the exact circumstances of the first infection have not been determined, but evolutionary and comparative analysis suggests a likely transmission from bats to pangolins to human ^2, 3^. Although the rate of mutation in the viral genome is believed to be slower than other viral pathogens, several variants of concern (VoC) have become dominant strains in different locations around the world such as the U.K. variant B.1.1.7 (Alpha, α), the South African variant B.1.135 (Beta, β), the Brazilian/Japanese variant P.1 (Gamma, γ), the Indian variant B.1.617.2 (Delta, δ), and the Californian variant B.1.427/B.1.429 (Epsilon, ε)^4^ {WHO EPI-WIN, Update 60 https://www.who.int/teams/risk-communication/epi-win-updates}. VoC are problematic because contain mutations in the viral spike (S) glycoprotein that enhance affinity to the host viral receptor human angiotensin converting enzyme 2 (hACE2) ^5, 6^. In addition, neutralizing antibodies (NAbs) produced by the immune response after natural infection or through vaccination have been shown to be less effective against VoC containing mutations in key regions of the S targeted by the NAbs ^7, 8^. Hence the need of rapid and accurate screen procedures to identify existing, or novel, drugs against newly emerging SARS-CoV-2 VoC.

Drug development for specific antiviral agents that combat COVID-19 is still an unmet urgent task. Several safe and effective vaccines have been approved and large-scale vaccination campaigns have already taken place since early 2021 ^9^. However, there will be still need for a multi-pronged approach to manage COVID-19 cases as highlighted by the situation with influenza virus that is responsible for a significant number of yearly infections associated with significant morbidity and mortality despite safe and effective vaccines being on place. Moreover, as long as there are large numbers of infections anywhere in the world, the risk for another resurgence of cases and emergence of virulent SARS-CoV-2 VoC remains high.

Remdesivir, an RNA dependent RNA polymerase (RdRP) inhibitor, is currently the only drug approved for the treatment of hospitalized COVID-19 patients ^10^. One challenge for remdesivir is that it needs to be administered intravenously ^11^. In addition, the therapeutic efficacy of remdesivir is also in question ^12 13 14^. Therefore, effective antiviral agents are urgently needed for treatment of COVID-19 patients.

Optimized compound screening assays are critically important for early drug discovery and development. Currently, two main types of assays are being use for compound screens in the COVID-19 drug development area of research. One is a mechanistic-based approach directly targeting SARS-CoV-2 proteins including 3CLpro ^15^, PLpro ^16^, RdRP ^17^, and S-hACE2 binding ^18^. Another approach is phenotypic screens using cell-based functional assays such as viral entry ^19, 20^, viral replication (replicon) ^21^, and live virus infection. The live SARS-CoV-2 assay is an important tool for compound screening and evaluation of efficacy before advancing forward to *in vivo* model efficacy studies. To date, assays including SARS-CoV-2 cytopathic effect (CPE) ^22^, RT-PCR, and antigen immunostaining, among others, have been used in compound screens and evaluation of drug efficacy ^23^. While the RT-PCR assay does not have enough screening throughput, the SARS-CoV-2 CPE assay may not identify all of the active compounds that inhibit SARS-CoV-2 replication in the screen because CPE is cell type-dependent and is an indirect readout of viral infection.

SARS-CoV-2 NP is a structural protein that binds the positive strand viral genome within the nucleocapsid core ^24, 25^. We developed an AlphaLISA assay for the measurement of SARS-CoV‑2 NP as an indicator of virus replication in infected cells ^26^. We used the NP as a target viral to develop a high-throughput screening (HTS) assay that could be used in a biosafety level 3 containment facility to identify promising antiviral drugs targeting SARS-CoV-2 multiplication ^27^. Targeting the more conserved and highly expressed NP instead of the S would prevent detection failures due to newly emerging SARS-CoV-2 VoC with mutations S mutations and increase sensitivity, respectively.

We have developed a homogeneous time resolved fluorescence (HTRF) assay that relies on the time-resolved fluorescence resonance energy transfer (TR-FRET) between a donor and acceptor fluorophore when they are in close proximity and the proper orientation relative to each other ^28, 29^. To specifically detect SARS-CoV-2 NP, we conjugated two NP-specific MAbs to either a donor or acceptor fluorophore that are brought into proximity when NP is present. We developed a microplate assay using recombinant NP added to cells, NP transfected cells, and both tissue culture supernatants (TCS) and cell lysates from infected cells. The homogeneous format means that the whole well fluorescence emission ratio between the acceptor and donor fluorophores is read without washing using a plate reader with a TR-FRET module.

The assay we have developed involves a single step, it is easy to use and scalable to HTS to identify SARS-CoV-2 antivirals. The assay can identify drugs targeting any of the steps of SARS-CoV-2 life cycle, hence enabling screening efforts to identify drugs with activity against emerging VoC.

## Methods

### Reagents

Vero E6 cells (CRL-1586, Resource Research Identifier (RRID): CVCL0574) were purchased from American Tissue Type Collection. The following items were purchased from Corning TM: EMEM (10–009-CV), HI FBS (35-016-VC, and 0.25% Trypsin (25053CI). Pen/Strep (15140–122) was purchased from Gibco. PBS (SH30256FS) was purchased from HyClone. The following items were purchased from Greiner Bio-One: white 384 well plate (781073), white half area 96-well plate (675083), white 96-well plate (655083). The custom labeling of MAbs was performed by Columbia Biosciences. Rabbit anti-NP SARS-CoV-2 antibody R001 (40143-R001, RRID Number: AB_2827974), R004 (40143-R004, RRID Number: AB_2827975), R019 (40143-R019, RRID Number: AB_2827973), and R040 (40143-R040, RRID Number: AB_2827976) was purchased from Sino Biological and mouse anti-NP MAb 1C7C7 was provided by Dr. Tomas Moran and purchased from Leinco (LT7000). The following items were purchased from ThermoFisher: Goat anti-mouse HRP (A16072), goat-anti rabbit HRP (A16104), West Femto substrate (34094), Lipofecatmine 2000 (11668019), Optimem I Reduced Serum Media (31985070). Triton X-100 (100x) was purchased from Sigma Aldrich. cOmplete™ ULTRA protease inhibitor (PI) (05892791001) was purchased from Roche.

### Antibody matrixing

Four rabbit MAbs and one mouse Mab specific for SARS-CoV-2 NP were labeled with Europium (donor Ab) or DyLight650^®^ (acceptor Ab). IgG antibodies R001, R004, R019, R040 were raised against recombinant SARS-CoV nucleocapsid phosphoprotein (NP (Sino Biological, 0143-V08B) and expressed in HEK293 cells. Mouse MAb 1C7 (1C7C7, IgG2a) expressed in HEK293 using recombinant NP as the immunogen. The labeled MAbs were then tested in HTRF assay, in triplicate wells, using a cross-matrix assay format. The data is presented as a median TR-FRET ratio (acceptor fluorescence/donor fluorescence X 10,000).

### Western blot

After SDS-PAGE and Western blotting the membranes were probed with anti-NP MAbs R001 and 1C7 at 1:1000 diluted in Superblock Buffer (ThermoFisher). Then antigen-antibody complexes were detected using appropriate anti species HRP-conjugates and West Femto substrate.

### Vero E6 Cell Culture

African green monkey kidney epithelial cells (Vero E6; CRL-1586) were grown and maintained in Dulbecco’s modified Eagle’s medium (DMEM) supplemented with 10% fetal bovine serum (FBS) and 1% PSG (100 U/ml penicillin, 100 μg/ml streptomycin, and 2 mM l-glutamine) at 37°C with 5% CO2.

### Assay condition optimization using recombinant NP

The best pair donor and acceptor, R001-Eu (donor) and 1C7-DL650 (acceptor), was used to evaluate assay performance in cell culture media with and without cells. Recombinant untagged NP starting at 1500 ng/mL was serially diluted two-fold 11 times to produce the standard curve. NP was diluted in media and 20 μL was added to the wells followed by 10 μL of 3x HTRF donor and acceptor diluted in 1.5% Triton X-100 and PI in the ratios indicated in Figure 2 for a total assay volume of 30 μL. In another 384-well plate, 15 μL of Vero E6 cells was plated and incubated overnight (O/N) prior to addition of 5 μL of 4x NP in media and 10 μL of 3x HTRF reagents in Triton X-100 with PI for a total assay volume of 30 μL. Plates were incubated with reagents for either 1h at RT or O/N at 4 °C. The TR-FRET ratio was measured at the end of each incubation time. Briefly, the 337 nm/620 nm excitation/emission and 337nm/665nm excitation/emission were measured using the dual emission HTRF optical module on the Pherastar (BMG Labtech). The ratio of emissions (665 nm/620 nm) was calculated and multiplied by 10000.

**Figure 1.**
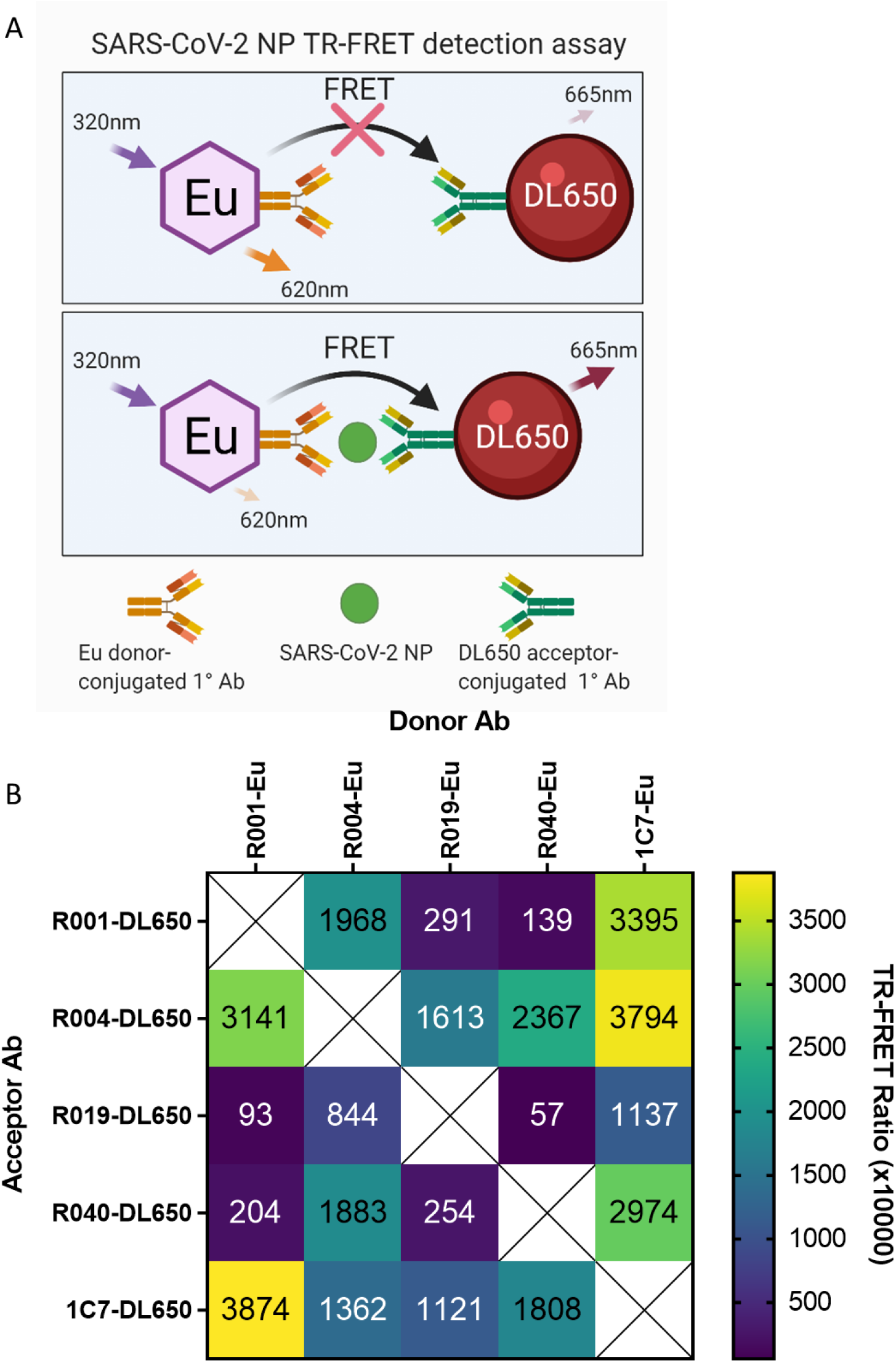
Selection of optimal donor/acceptor antibody pair. **(A)** Illustration of the HTRF assay for SARS-CoV-2 NP showing the Eu donor-conjugated primary MAb and DL650 acceptor-conjugated primary MAb detecting SARS-CoV-2 and enabling FRET. **(B)** Five MAbs specific for SARS-CoV-2 NP were labeled with Europium (donor Ab) or DyLight650^®^ (acceptor Ab). The labeled MAbs were then tested in HTRF assay, in triplicate wells using a cross-matrix assay format. The presented data is a median TR-FRET counts (acceptor fluorescence/donor fluorescence X 10,000). The MAb pair (R001-Eu/1C7-DyLight650) showed highest specific signal and was selected for further assay development.

**Figure 2.**
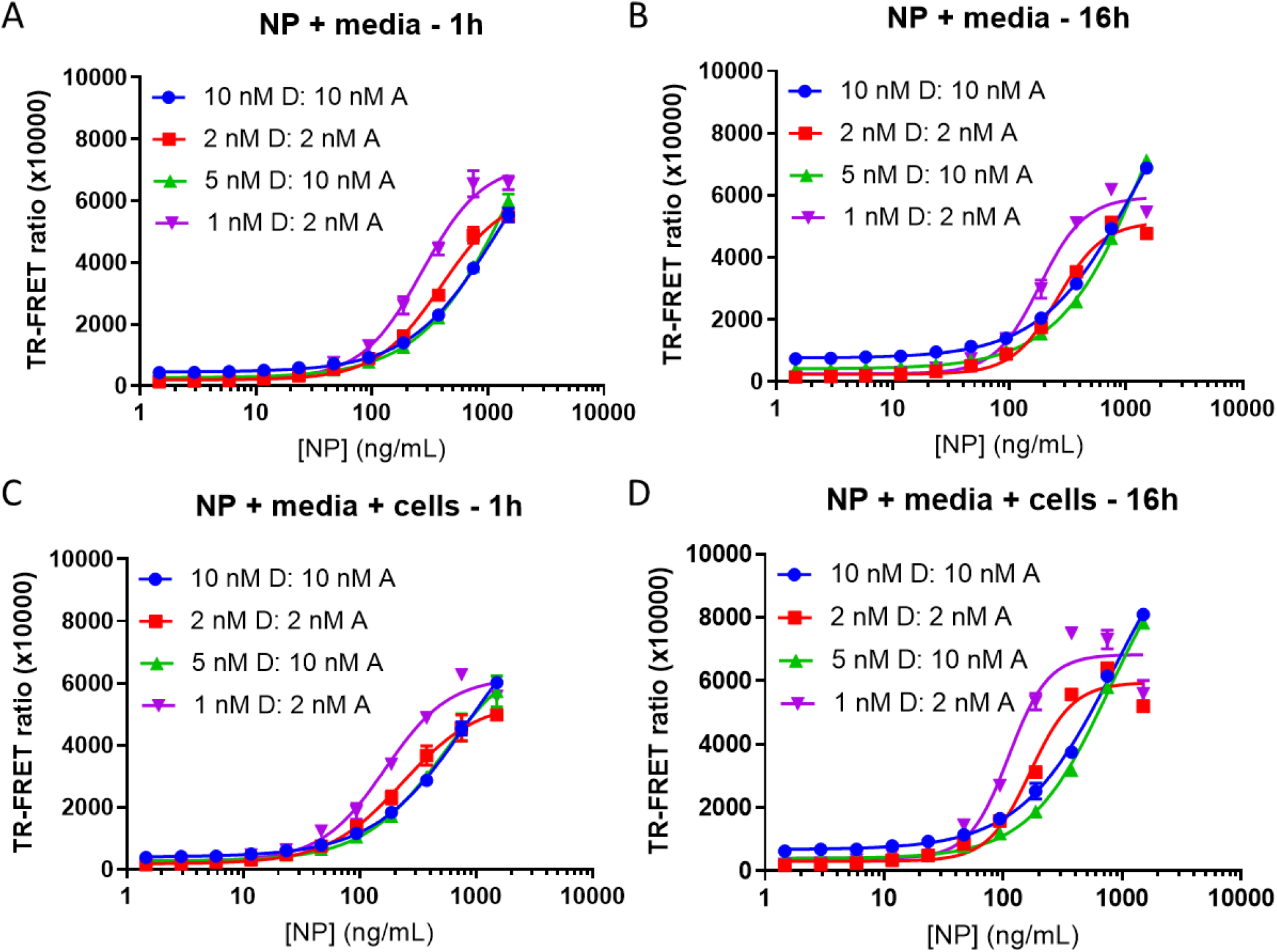
Optimization of donor to acceptor ratio, concentration, and incubation time using media or Vero E6 cells. TR-FRET ratio signal (acceptor/donor) detection using R001-Eu and 1C7-DL650 as the donor (D) and acceptor (A) pair respectively. SARS-CoV-2 NP was used to generate the 11-point standard curve starting from 1500 ng/mL serially diluted 1:2. HTRF reagents were incubated with NP in cell culture media only **(A and B)** or in media with 5000 Vero E6 cells **(C and D)** for either 1h at RT or O/N at 4°C. N = 3 wells per condition in a 384 well plate. Error bars indicate S.D.

### Preparation of Viral Lysates and Tissue Culture Supernatant (TCS) Production

Vero-E6 cells were plated in 12-well plates at 450 000 cells/well in 1.25 mL of growth media. Cells were incubated for 24h at 37 °C. A 250 μL aliquot of SARS-CoV-2 [USA-WA1/2020 strain (Gen Bank: MN985325.1)] was used at an MOI of 0.05. Cells were inoculated for 45 min at 37 °C. Supernatant was collected and pooled for 0h TCS (Mock). A final concentration of 0.5% Triton-X 100 and 1× protease inhibitor (PI) was used for all samples. TCS samples were stored at −20 °C until needed. Next, 1.5 mL of fresh media was added to each well, and cells were incubated for 24h and 48h at 37 °C. Six wells were harvested at each time point. Supernatant was collected from 6 wells each at 24 and 48 h. Samples were pooled for Mock, 24h or 48h TCS. Samples were stored at −20 °C until needed. Lysate was collected from 6 wells each at 24 and 48 h. Cells were rinsed once with ice-cold PBS, and 250 μL/well of cell lysis buffer + PI was added to lyse cells. Protease inhibitor was added to the lysis buffer before lysing cells. Cells were scraped to the bottom of the well, and pooled lysates were collected into 1.5 mL tubes on ice. Tubes were vortexed for 10 s and placed on ice for 10 min. This was repeated three times. Lysates were spun down for 30 min at 13200 rpm at 4 °C and stored at −20 °C until further use. Estimates of viral NP concentrations in infected TCS and cell lysate were conducted using standard curve interpolation within the linear range.

### Testing TCS and cell lysates from infected Vero E6 cells

This TR-FRET NP assay was evaluated for performance using Vero E6 TCS and lysate in 384 well plates. Lysates and TCS were diluted in PBS + Triton-X 100 + protease inhibitors 1:15 (1×) followed by 3-fold dilutions. Plates were incubated for either 1h at RT or O/N at 4 °C. The TR-FRET ratio was measured at the end of each incubation time.

### Evaluating 384-Well plate assay robustness

Whole-plate statistics were calculated using recombinant NP at a concentration of 500 ng/mL in Vero E6 cells. 15 μL of 5000 Vero E6 cells was seeded into 384 well plates and incubated O/N. Reagents were added using a fully automated Thermo Fisher Combi Multidrop liquid dispenser. 5 uL of cell culture media with or without NP was added to columns 3–24 and columns 1–2, respectively. Next, 10 μL of 3x HTRF reagents in Triton X-100 with PI for a total assay volume of 30 μL. The TR-FRET ratio was measured at the end of each incubation time. Z-factor, CV, and signal to background (S/B) was calculated for the entire plate.

### Transfection of NP into Vero E6 cells

DNA coding sequences corresponding to SARS-CoV-2 NP were amplified from genomic RNA from SARS-CoV-2, Isolate USA-WA1/2020 (BEI Resources, Catalog No. NR-52285) using primer sets 5SacINP (5’- CACTGAGCTCATGTTTGTTTTTCTTGTTTTATTG-3’) and 3SmaINP (3’- CACTCCCGGGTGTGTAATGTAATTTGACTCCTTT-5’). PCR fragments were digested with SacI and SmaI and cloned into SacI- and SmaI-digested pCAGGS plasmid containing a C-terminal HA epitope tag (pCAGGS NP-HA)

20 uL of 15,000 Vero E6 cells per well was seeded into half-area 96-well plates and incubated O/N at 37°C, 5% CO_2_. A plasmid encoding for SARS-CoV-2 NP (pCAGGS NP-HA) was transfected into the cells using Lipofectamine 2000 according to manufacturer’s recommendations (starting concentration 100 ng of plasmid and two-fold serial dilutions). Cells were incubated with plasmid and Lipofectamine 2000 for 4-6h before replacing media with DMEM containing 2% FBS. Cells were incubated O/N at 37°C, 5% CO_2_ to express NP in a volume of 20 μL. After 24 h, 10 μL of 3x HTRF reagents diluted in 1.5% Triton X-100 with PI were added to cells and incubated for 1h at RT or O/N at 4 °C. The TR-FRET ratio was measured at the end of each incubation time.

### Live virus testing in Vero E6 cells

A reverse time course was used to infect cells. Briefly, 20 μL of Vero E6 cells was seeded into half-area 96-well plates at 8500, 6000, and 4500 cells/well for the 48h, 24h, and 12h infection durations respectively and incubated O/N at 37°C, 5% CO_2_. 4500 cells/well were also used for the 0h infection duration and the standard curve. Media was removed the next day and 20 μL/well of 3-fold serial dilutions (starting MOI 0.25) of SARS-CoV-2 USA-WA1/2020 (BEI Resources, NR-52281). Upon completion of the reverse time course 10 μL of 3x HTRF reagents diluted in 1.5% Triton X-100 with PI was added to cells and incubated for 1h at RT or O/N at 4 °C. The TR-FRET ratio was measured at the end of each incubation time.

### Demonstrating antiviral efficacy using remdesivir

20 μL of Vero E6 cells was seeded into half-area 96-well plates at 8500 cells/well and incubated O/N at 37°C, 5% CO_2_. Cells were infected with 10 μL of SARS-CoV-2 USA-WA1/2020 in infection medium (DMEM with 2% FBS) at an MOI of 0.009, cells were simultaneously treated with 10 μL of 3-fold serial dilutions (starting concentration of 100 μM) of remdesivir or 0.1% dimethyl sulfoxide (DMSO) as a vehicle control. After 24h, 10 μL of 3x HTRF reagents diluted in 1.5% Triton X-100 with PI was added to cells and incubated for 1h at RT or O/N at 4 °C. The TR-FRET ratio was measured at the end of each incubation time.

### Detecting variants of concern

A reverse time course was used to infect cells. 20 μL of Vero E6 cells were seeded into half-area 96-well plates, at 8500 cells/well for 24 h and 6000 cells/well for 12 h, 8 h, 4 h, 0 h infection durations as well as the standard curve, and incubated O/N at 37°C, 5% CO_2_. Media was removed the next day and cells were infected with 20 μL/well of 3-fold serial dilutions (starting MOI 0.25) of SARS-CoV-2 isolates USA-WA1/2020, SARS-CoV-2/human/USA/CA-UCSF-0001C/2020 (UCSF, a gift from Dr. Charles Chiu, hCoV-19/Japan/TY7-501/2021 (BEI Resources, NR-54981), or hCoV-19/South Africa/KRISP-EC-K005321/2020 (BEI Resources, NR-54008) in infection medium (DMEM with 2% FBS) for each indicated time point. 10 μL of 3x HTRF reagents diluted in 1.5% Triton X-100 with PI were added to cells and incubated for 1h at RT or O/N at 4 °C. The TR-FRET ratio was measured at the end of each incubation time.

### NP amino acid sequence alignment

Nucleotide sequences of viral strains, SARS-CoV-2 isolate USA-WA1/2020 (accession number EPI_ISL_404895), Beta (EPI_ISL_678625), Gamma (EPI_ISL_833366) and Epsilon (EPI_ISL_2712562) were downloaded from GISAID ^30^. EPI_ISL_2712562 only used for alignment, contains same mutations in NP as SARS-CoV-2/human/USA/CA-UCSF-0001C/2020 used in this study. Nucleotide sequences were translated to amino acid sequences using ExPASY online software ^31^ and the codon frame that contained the full NP was copied to ClustalO (1.2.4)^32^ for multiple sequence alignment. Portions of the sequences that contained mutations were extracted and compared.

### Statistical analysis

Nonlinear regression was used to generate the curve-fits. S/B calculations were made using the ratio of the TR-FRET signal at time X and the signal at time 0. For TCS and lysate NP concentration interpolation, an asymmetric sigmoidal 5PL model was used. Graphpad Prism V9.1.0 was used to generate all graphs. Biorender was used to generate the illustration in Figure 1A.

## Results

### Selection of donor and acceptor MAb pairs for the TR-FRET NP assay

We sought to develop a precise, reproducible, scalable, and easy to use assay to detect infection and replication of SARS-CoV-2 under conditions compatible with HTS. We selected SARS-CoV-2 NP as the target viral protein to be a direct readout of viral infection and replication. We chose a FRET-based strategy using the popular HTRF system consisting of a donor and acceptor fluorophore conjugated to two SARS-CoV-2 NP MAbs, as these detection reagents offer high sensitive and dynamic range to make HTS possible. The detection system consisted of the lanthanide fluorophore europium cryptate (Eu donor) conjugated to one NP MAb, and the acceptor fluorophore DyLight 650 (DL650 acceptor) conjugated to a second NP MAb (Figure 1A). In the absence of SARS-CoV-2 NP, the distance and orientation of the donor and acceptor are incompatible with energy transfer, whereas in the presence of NP the proximity of donor and acceptor is close enough to allow for FRET to occur. Laser excitation of Eu at 320 nm induces an emission at 620 nm. With energy transfer from the Eu donor, the DL650 acceptor produces an emission at 665 nm. The ratio of acceptor to donor emission intensity (665 nm/620 nm) upon excitation at 320 nM is calculated as the TR-FRET signal that correlates positively with the amount of NP present. In the absence of NP, the ratio is smaller and any signal present is considered background signal produced by random interactions between donor and acceptor.

We tested several combinations of primary SARS-CoV-2 NP MAbs and the greatest TR-FRET signal was produced by R001 coupled to the Eu donor and 1C7 coupled to the DL650 acceptor (Figure 1B). This combination was selected for further development. Both MAbs detected SARS-CoV-2 NP as shown by Western Blot (Supplementary Figure 1).

### Evaluation of assay performance for detection of recombinant SARS-CoV-2 NP

We evaluated cell culture media, the presence of cells, reagent incubation time, the ratio of donor to acceptor, and the concentration of reagents on the TR-FRET ratio and signal to background ratio (S/B) in the presence of recombinant NP. In media alone with a 1h incubation of reagents, a concentration of 1 nM donor and 2 nM acceptor produced the greatest TR-FRET ratio, was more sensitive at lower concentrations of NP, produced the largest S/B, but started to plateau at the highest concentrations of NP. The maximum ratio achieved at the highest concentration of NP (1500 ng/mL) was comparable between all reagent combinations (Figure 2A).

After O/N incubation, a hook effect appears with the lower concentration of reagents at the highest concentration of NP, suggesting an upper limit of detection between 750 ng/mL and 1500 ng/mL of NP. The hook effect is defined as a signal decrease due to saturating antigen concentrations that prevent binding of both donor and acceptor linked antibodies to the same antigen molecule. 1 nM donor and 2 nM acceptor was saturated at 1500 ng/mL of NP, but exhibited high S/B (Figure 2B, Supplementary Figure 2). In contrast with the low concentration combinations, the high concentrations of reagents did not saturate, suggesting a higher upper limit of detection than 1500 ng/mL of NP.

In the presence of cells, the S/B was elevated (Figure 2C,D, Supplementary Figure 2) compared to media alone. The overall greatest S/B was achieved with 1 nM donor and 2 nM acceptor (Supplementary Figure 2), although the hook effect was more apparent for the low concentrations of reagent. The balance of sensitivity (lower limit) and dynamic range (upper limit) with S/B is an important consideration when choosing the conditions that are optimal for viral detection. The combination of 1 nM donor and 2 nM acceptor produces acceptable ratios with good sensitivity while reducing reagent consumption, but has a reduced upper limit of detection.

### Evaluation of NP detection SARS-CoV-2 infected tissue culture supernatant (TCS) and cell lysates

We next evaluated the assay performance in TCS and whole cell lysates of Vero E6 cells infected with SARS-CoV-2 USA-WA1/2020 strain for 24h or 48h using mock-infected cells as negative control. Using 1 nM donor and 2 nM acceptor, the TR-FRET ratio was greatest with cell lysate, but a low level signal was detected in TCS as well (Figure 3A-D, Supplementary Figure 3). Interestingly, the sensitivity of NP detection in these samples increased with O/N incubation as evidenced by the increased TR-FRET ratios and hook effect at lower dilutions of lysate (Figure 3C,D, Supplementary Figure 3). Using standard curve interpolation of the O/N incubation TR-FRET values, NP concentration of 70.9 ± 12.3 ng/mL and 1050 ± 12.3 ng/mL in TCS after 24h and 48h were observed, respectively. The lysates exhibited larger NP concentrations of 16380 ± 1290 ng/mL and 59520 ± 957 ng/mL after 24h and 48h infection, respectively (Figure 3D). The data indicated that both recombinant and nascent NP produced in cells after viral infection could be detected by our TR-FRET assay.

**Figure 3.**
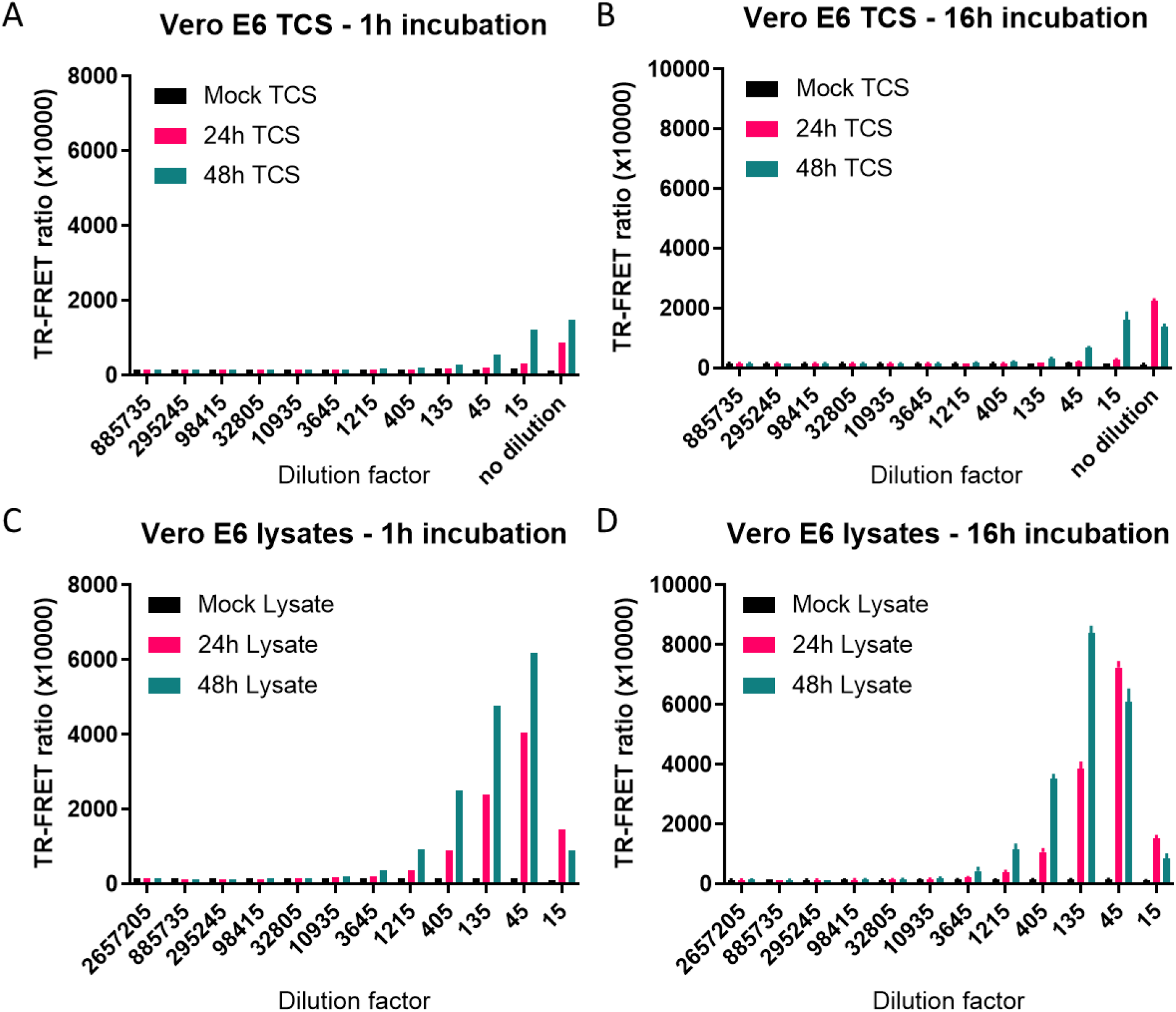
Detecting SARS-CoV-2 NP from Vero E6 TCS and cell lysate. TCS and cell lysate were collected from Vero E6 cells after infection with SARS-CoV-2 USA-WA1/2020 strain for 24h or 48h. TR-FRET ratio from TCS **(A and C)** and cell lysates **(B and D)** after incubating with reagents for 1h at RT or O/N at 4°C. TCS was diluted 1:3 and cell lysate was first diluted 15-fold followed by 1:3 dilutions. N = 3 wells in a 384 well plate. Error bars indicate S.D.

### Assessing HTS readiness of the assay

To evaluate the suitability of the assay for HTS, we seeded Vero E6 cells into a 384-well plates using an automated liquid handler (ThermoFisher Multidrop Combi) and dispensed 500 ng/mL of recombinant NP into columns 2-24 (Figure 4). Columns 1-2 were used as a negative control. After addition of 1 nM donor and 2 nM acceptor, the plate was read at 1h and after O/N incubation at 4°C. The assay performance was excellent with coefficients of variation (CV) below 5%, an S/B above 36, and a Z-factor at or above 0.90. O/N incubation increased S/B to 50.7, compared to 36.6 after 1h incubation, and also decreased the CV, and improved the Z-factor.

**Figure 4.**
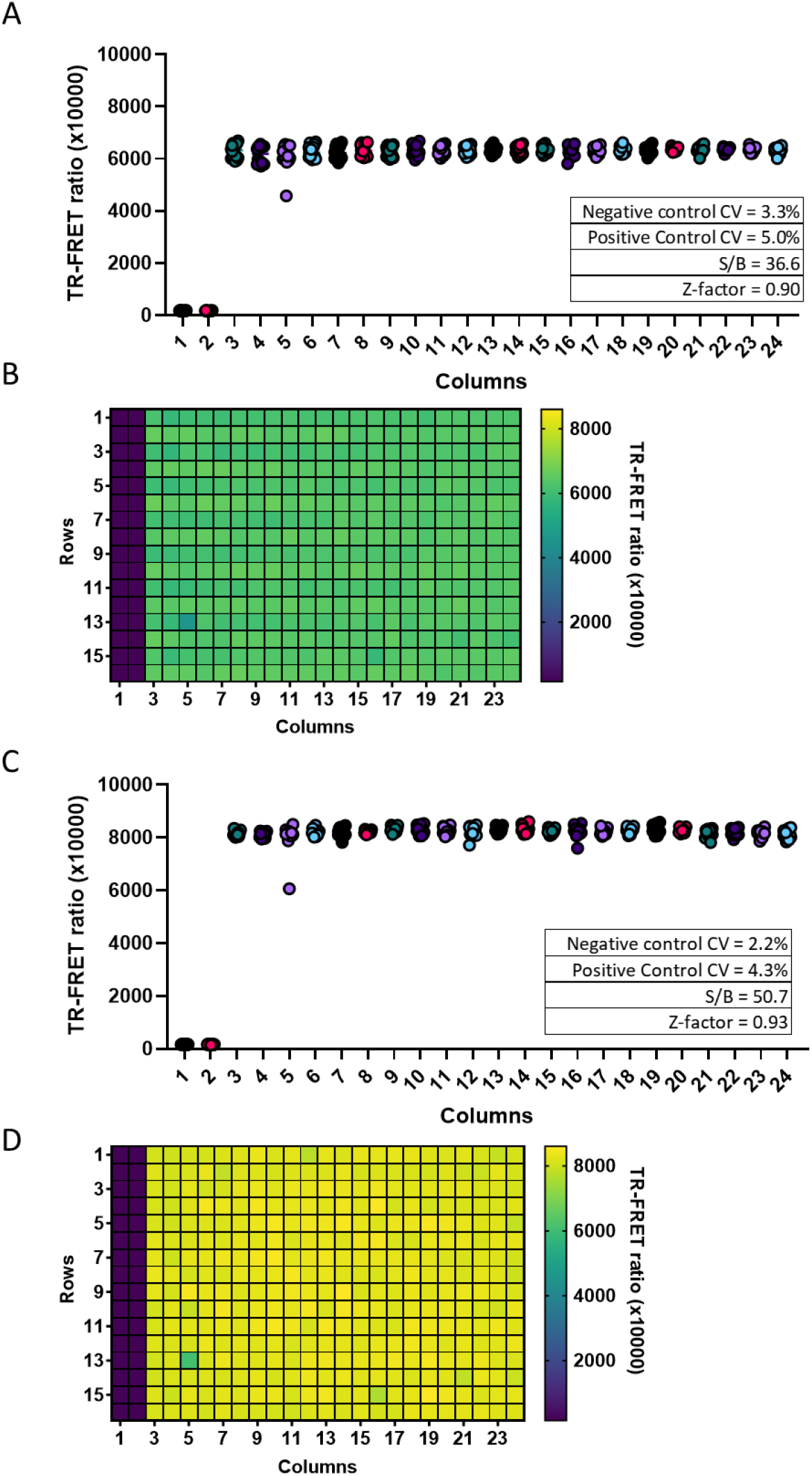
384 well plate statistics using spiked NP in Vero E6 cells. Vero E6 cells grown in a 384 well plate at 5000 cells per well were spiked with 0 ng/mL or 500 ng/mL of SARS-CoV-2 NP and incubated with HTRF reagents for 1h at RT **(A and B)** or **O/N** at 4°C **(C and D)**. Columns 1 and 2 contained no NP and acted as a simulated positive control for uninfected cells. Columns 3 to 24 contained 500 ng/mL of SARS-CoV-2 NP. Each column contains 16 wells. Each point represents 1 well. Inset, calculated plate statistics

### SARS-CoV-2 NP TR-FRET assay detects live virus infection in Vero E6 cells

To assess whether our assay reagents were able to access viral NP produced in SARS-CoV-2 infected live cells, we first transfected Vero E6 cells with different amounts of an NP expression plasmid for 24h (Supplementary Figure 4), followed by addition of HTRF reagents at high and low concentrations and incubation for either 1h or O/N at 4°C. After 1h, the low concentration was more sensitive, and the high concentration produced larger TR-FRET ratio values with higher amounts of plasmid. O/N incubation shifted the curves leftward, increasing assay sensitivity and S/B.

We then tested this NP TR-FRET assay using SARS-CoV-2 -infected Vero-6 cells (Figure 5A). Production of SARS-CoV-2 NP in infected cells can be experimentally modulated by adjusting the multiplicity of infection (MOI) and infection time. To determine optimal assay conditions for the detection of SARS-CoV-2 NP in live cells, we seeded Vero E6 cells in 96-well plates and infected them with the SARS-CoV-2 USA-WA1/2020 for 48h, 24, 12, and 0h (Figure 5A). We used 1:2 ratios of donor and acceptor and tested high (5 nM D: 10 nM A) and low concentrations of reagents (1 nM D: 2 nM A) with 1h and O/N incubation. At 0h, an MOI between 0.0003 and 0.0001 started producing detectable signal above background that was caused by the NP present in the virus inoculum (Figure 5B). The low concentrations of reagents were more sensitive at lower MOIs, but the high concentrations produced the larger TR-FRET signal at higher MOI. The high concentrations of reagents exhibited a hook effect at higher MOI than the low concentrations of reagents. At 12h, the MOI-response curves shifted leftward towards lower MOI, suggesting the virus was replicating and producing more NP (Figure 5C). At 24h, the low concentrations of reagents could no longer accommodate the high levels of NP produced at high MOI, and low MOI had greater variability (Figure 5D). The high concentration of reagents more accurately detected the NP produced in viral infected Vero E6 cells and the hook effect was observed at MOI greater than 0.003. At 48h, the viral NP concentration was saturated at all MOIs and could not be accurately detected (Figure 5E). The standard curve was added for reference (Figure 5F). Thus, an MOI of 0.009 for SARS-CoV-2 infection followed by a 24 h incubation and 5 nM donor with 10 nM acceptor was chosen as an optimized assay condition for following experiments.

**Figure 5.**
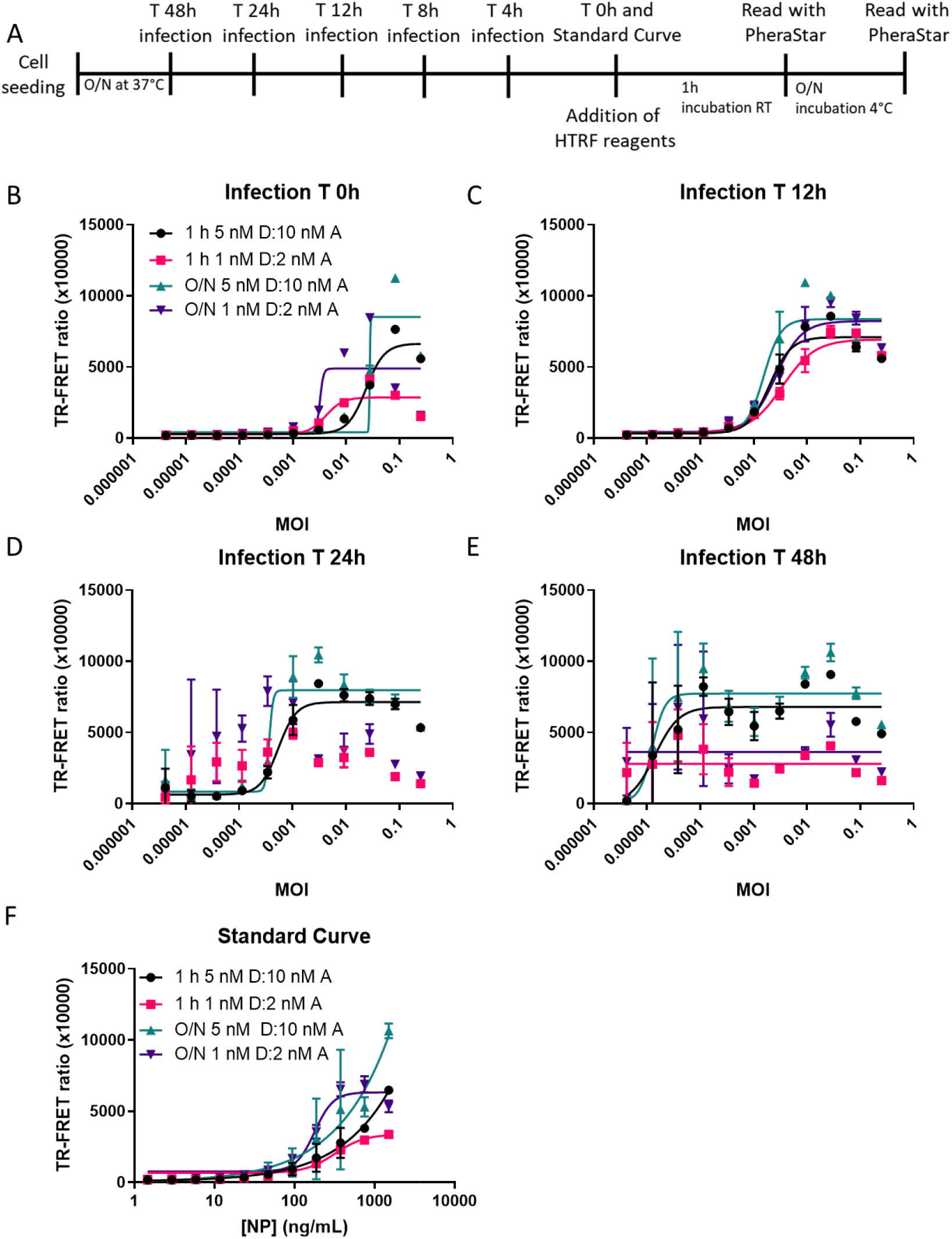
HTRF assay detects NP produced by live virus infection of Vero E6. The TR-FRET ratio for SARS-CoV-2 NP detected in Vero E6 infected with the SARS-CoV-2 USA-WA1/2020 strain at different MOI starting at 0.25 and diluted 1:3 based on a reverse time-course protocol (**A**) for 0 h (**B**), 12 h (**C**), 24 h (**D**), or 48 h (**E**). The standard curve **(F)** with assay conditions equal to the 48h time-point. Plates were incubated with NP HTRF reagents for 1h at RT or O/N at 4°C. N=3 wells in a half-area 96-well plate. Error bars indicate S.D.

### Use of the SARS-CoV-2 NP TR-FRET assay to assess the effect of Remdesivir on NP levels in SARS-CoV-2 infected Vero E6 cells

To assess the utility of our assay to determine the potencies of antiviral drugs, we examined the effect of Remdesivir on NP levels in SARS-CoV-2 infected Vero E6 cells. For this, we treated SARS-CoV-2 USA-WA1/2020-infected (MOI 0.009) Vero E6 cells with Remdesivir at the time of infection (Figure 6A). After 24h, 5 nM donor and 10 nM acceptor were added to the cells and the TR-FRET ratio was measured after 1h and O/N incubation. We observed a half maximal inhibitory concentration (IC_50_) of 9.3 μM and 9.5 μM with 1h and O/N incubation, respectively (Figure 6B). The standard curve was added for reference (Figure 6C). Remdesivir completely inhibited SARS-CoV-2 replication as determined by the very low TR-FRET ratio at the highest concentration of drug. Residual NP from the initial inoculation likely accounted for low amount of signal seen at the highest concentration of drug.

**Figure 6.**
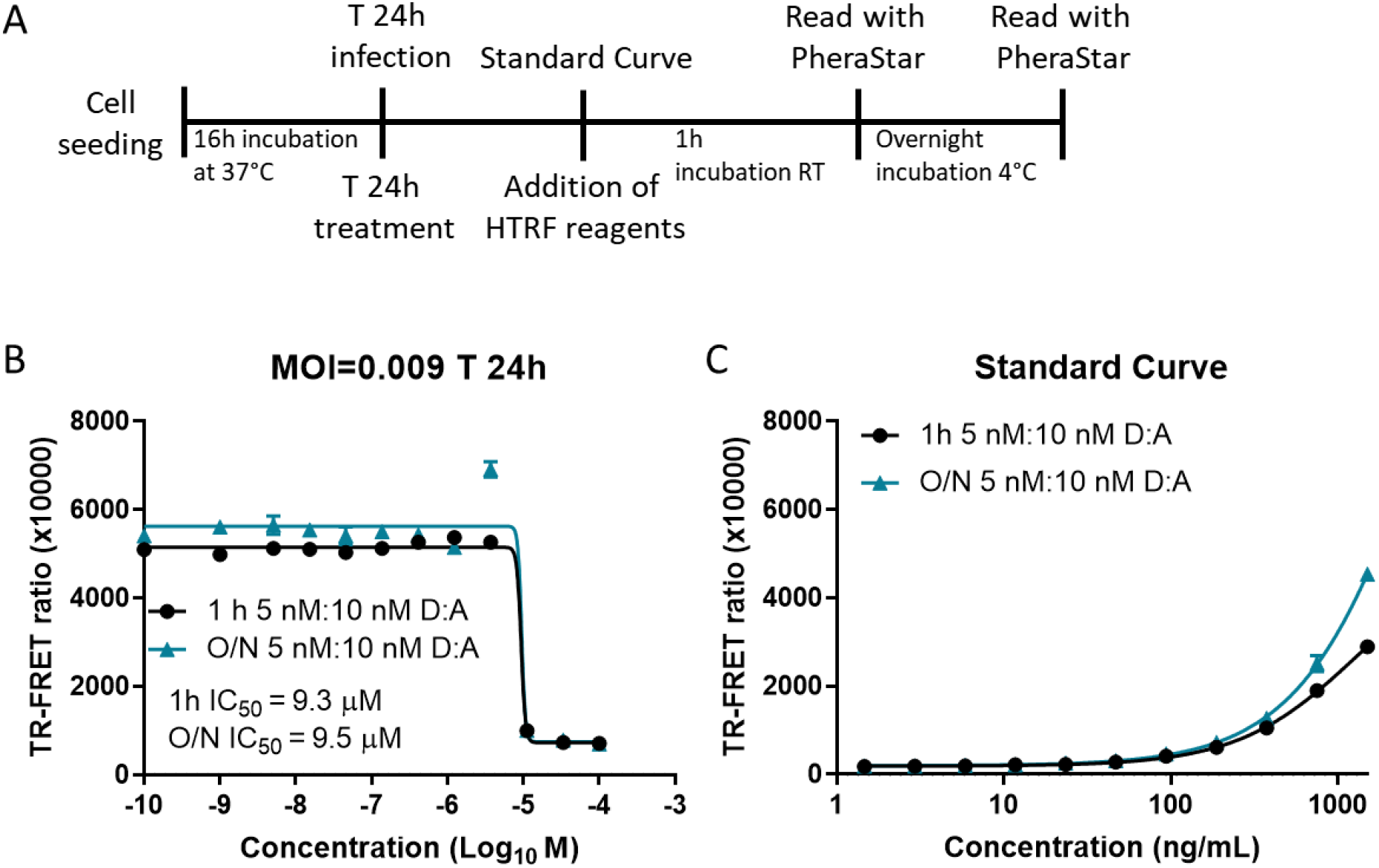
NP HTRF assay confirmation of remdesivir inhibition of SARS-CoV-2 replication. The TR-FRET ratio for SARS-CoV-2 NP in Vero E6 infected with SARS-CoV-2 USA-WA1/2020 strain at MOI 0.009 based on protocol (**A**) for 24 h (**B**). Remdesivir was added at a starting concentration of 100 μM and serially diluted 1:3. The standard curve (**C**) using recombinant SARS-CoV-2 NP. Plates were incubated with NP HTRF reagents for 1h at RT or O/N at 4°C. N=3 wells in a half-area 96-well plate. Error bars indicate S.D.

### SARS-CoV-2 NP HTRF assay detects VoC

Mutations in response to selective pressures driving SARS-CoV-2 evolution are heavily weighted towards the viral S protein because of its importance in cell entry and evasion of immune responses. Given that our assay depended on the detection of NP, highly mutated NPs may prevent MAb binding and would jeopardize the use of the assay. To examine whether this represented a problem for the detection of SARS-CoV-2 VoC, we infected Vero E6 cells with the SARS-CoV-2 USA-WA1/2020, Beta, Gamma, and Epsilon variants, which contained mutations in the S glycoprotein and NP, for 24, 12, 8, 4, and 0 h (Figure 7A, Supplementary Figure 5). We aligned the NP amino acid sequences (GISAID database) from the viruses used in this study and found several mutations including P80R (Gamma), R203K (Gamma, Epislon), G204R (Gamma), T205I (Beta, Gamma), and M234I (Epsilon) (Supplementary Figure 5). At the end of the reverse timecourse, 5 nM donor and 10 nM acceptor were added to the wells and incubated O/N at 4^0^C. The TR-FRET readings suggested that the assay could detect the VoC with the S glycoprotein mutations and could accommodate differences in NP amino acid sequence as well. The increase in TR-FRET ratio was first observed at 8h and the curve was further shifted leftward at 12h and 24h, indicating increasing amounts of NP production with lower MOI (Figure 7B-F). The standard curve was added for reference (Figure 7G). Interestingly, SARS-CoV-2 USA-WA1/2020 had the largest TR-FRET ratio at early timepoints and at later timepoints exhibited the hook effect with high MOIs. The VoC exhibited low, but detectable TR-FRET ratios at early timepoints, and the hook effect was less apparent with the three VoC at high MOIs. Altogether, the results suggested that the assay could be used to detect SARS-CoV-2 VoCs and has the potential to be used in HTS to identify potent antiviral compounds and biologics against these and potentially future SARS-CoV-2 VoC.

**Figure 7.**
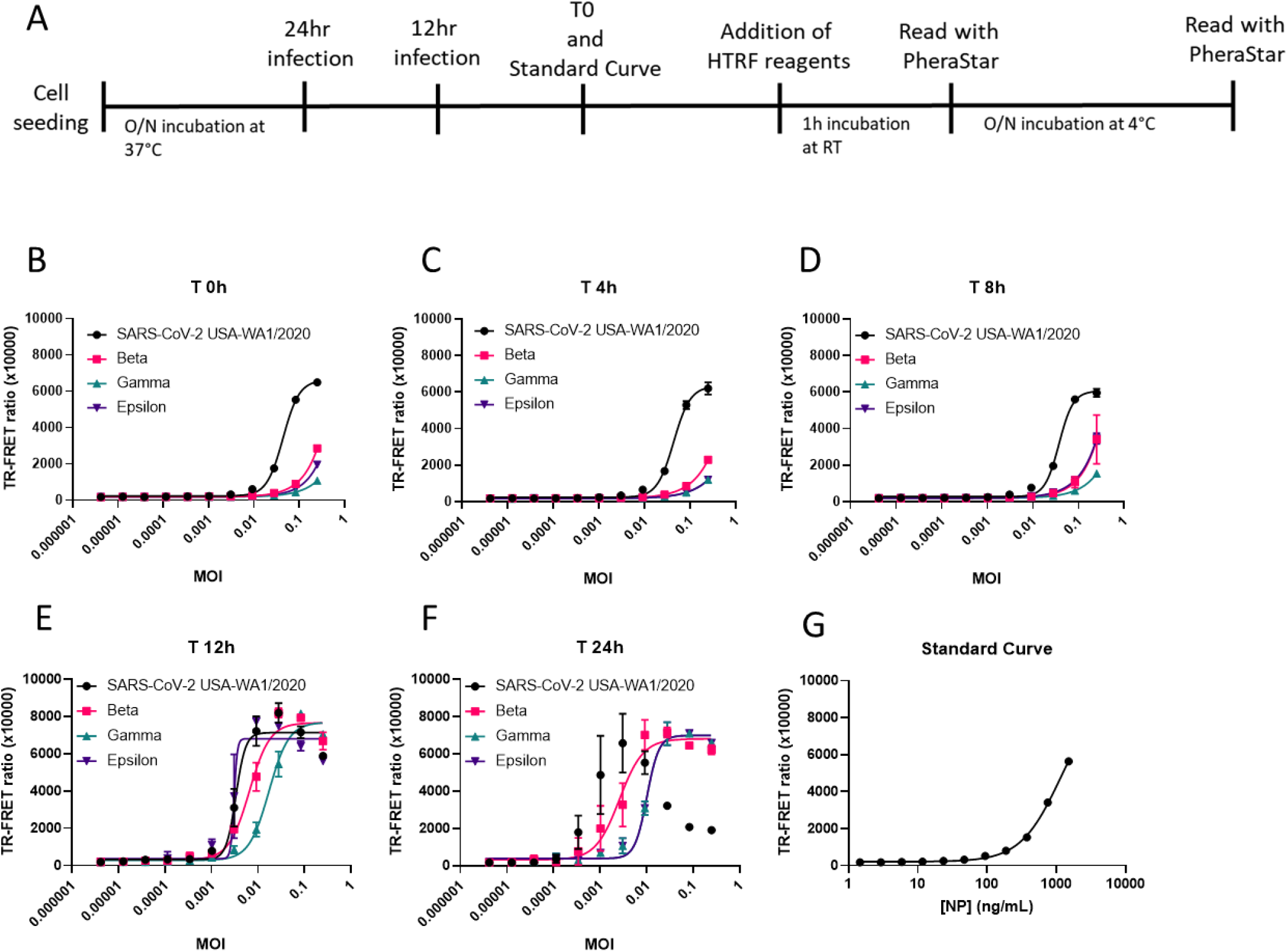
NP HTRF assay detects Beta, Gamma, and Epsilon VoC. The TR-FRET ratio for SARS-CoV-2 NP in Vero E6 infected with the USA-WA1/2020, Beta, Gamma, and Epsilon strains using protocol **(A)** at MOI starting at 0.25 and diluted 1:3 for 0 h (**B**), 4 h (**C**), 8 h (**D**), 12 h (**E**), 24 h (**F**). The standard curve **(G)** with assay conditions equal to the 24h time-point. Assay completed with a reverse time-course protocol; O/N incubation at 4°C with HTRF reagents (5 nM D/10 nM A). N=3 wells in a half-area 96-well plate. Error bars indicate S.D.

## Discussion

For early drug discovery and development, compound screening assays such as the one described herein are critical for lead identification and evaluation of molecules for further development. In this work, we demonstrated the robust and reliable detection of SARS-CoV-2 NP using the TR-FRET assay format in live virus-infected cells. This was further validated by the confirmation of efficacy of the well-characterized reference compound Remdesivir. We also demonstrated that this assay can also be used to measure virus replication and could be used in the future to evaluate activities of antiviral agents against emergent SARS-CoV-2 VoC.

This TR-FRET assay employs two specific MAbs to detect SARS-CoV-2 NP. Because the TR-FRET signals are produced when both MAbs bind to same NP, the assay specificity is usually high. The use of time-resolved fluorescence instead of flash fluorescence significantly increases assay sensitivity and reduces false positives in HTS due to compound autofluorescence. The ratiometric readout reduces well-to-well variations in assay plates that may be caused by different cell densities or reagent dispensing errors.

This assay development work reveals several key insights into the appropriate detection of SARS-CoV-2 NP produced during viral replication in host cells. One important aspect is the balance between sensitivity at the lower limit of detection, the dynamic range at the upper limit of detection, and the S/B that vary depending on the cell number, MOI, and incubation time after the virus infection. At very low MOI, the number of viral particles added to each well can be more variable than at high MOI where more virus particles per unit volume are evenly distribution to host cells. In this TR-FRET assay, signal positively correlates with MOI or incubation time after virus inoculation because of the expression of SARS-CoV-2 NP. At high MOI, the hook effect may become prominent because NP is expressed at higher levels. Because the concentrations of two labeled MAbs are fixed, the TR-FRET signal reaches a maximal value and then decreases with the further increases of MOI or incubation time due to high antigen (NP) concentrations that prevent binding of both donor and acceptor to the same antigen molecule. This unusual effect of reduced TR-FRET signals at higher MOI and longer incubation times (more viral replication cycles occur) is caused by the hook effect, but can be avoided with assay optimization that identifies an optimized assay condition. In this assay, an MOI of 0.009 and 24h infection time produced a clean MOI-FRET ratio curve when using the high 1:2 concentration of donor to acceptor. However, other concentrations and ratios may work better in different contexts such as different types of cells and different SARS-CoV-2 strains. For HTS, reagent conservation is key to keeping assay costs down and increasing the number of compounds to be screened. We demonstrated that relatively low concentrations of reagents produced robust assay signal for an HTS campaign.

The differences we observed in the detection of NP from the different VoC are interesting and could be due to several factors. First, the tropism of a virus for a particular cell line can be enhanced by adapting the virus to the cell line ^33^. Differences in the number of passages of different virus strains may cause the variations compared to the SARS-CoV-2 USA-WA1/2020 strain that we used in this study. Another possibility is that the NP produced by different VoC may have mutations that leads to different binding affinities of labeled MAbs used in this assay, causing changes in the FRET between the donor and acceptor ^34–36^. An analysis of the mutations suggests minimal changes to the NP for these mutants with one P80R mutation in the nucleotide binding region in the N-terminal domain of Gamma. The other mutations are all in disordered regions of the middle of the protein. Nonetheless, each SARS-CoV-2 VoC was detected and exhibited both a time-dependent and MOI-dependent increase in the TR-FRET ratio, enabling researchers to utilize this assay for the evaluation of antiviral agent efficacies against VoC, as well as HTS campaigns to identify new antiviral compounds. Compounds that are pan-active against SARS-CoV-2 VoC would be the most attractive candidates for further development.

In contrast to previously described CPE assays for SARS-CoV-2 infected cells, which depend on a cell-killing effect, the TR-FRET assay we described in the present paper measures NP produced in cells with active SARS-CoV-2 replication, without being dependent on cell death. Thus, this assay may be more biologically relevant because not all cell types are killed by SARS-CoV-2 ^22, 37^. The CPE may not be seen in all human cells *in vivo*. It should be noted that only a small portion of the hits found in an HTS using the TR-FRET based Zika virus non-structural protein 1 assay was confirmed by CPE assay (unpublished data). We believe that this TR-FRET SARS-CoV-2 NP assay will have a broad application in BSL-3 settings for lead compound identification in HTS and evaluations of antiviral therapeutics in emerging VoC.

## Abbreviations

TR-FRET: time resolved fluorescence resonance energy transfer
NP: nucleocapsid phosphoprotein
S: spike protein
VoC: variant of concern
NAbs: neutralizing antibodies
MAbs: monoclonal antibodies
MOI: multiplicity of infection
CPE: cytopathic effect
HTS: high-throughput screening
TCS: tissue culture supernatant

## Acknowledgements

We thank Dr. Charles Chiu, M.D./Ph.D., Director, UCSF-Abbott Viral Diagnostics and Discovery Center at UCSF School of Medicine for the isolate SARS-CoV-2/human/USA/CA-UCSF-0001C/2020. This research was supported in part by the Intramural Research Program of the National Center for Advancing Translational Sciences, NIH. We thank the development group at Columbia Biosciences for their assistance in reagent development.

## Author Contributions

Initial Concept – KG, CZC, LMS, WZ

Experimental – KG, DMV, BNT, JCT

Preparation of reagents – TM, KC, CY

Figure Preparation – KG, DMV

Manuscript writing – KG, DMV, WZ

Manuscript editing – KG, DMV, LMS, WZ

## Declaration of conflicts

The authors claim no conflicts of interest in the preparation of this manuscript.

## Notes

### Competing Interest Statement

The authors have declared no competing interest.

