## Supplementary Figures for "A SARS-CoV-2 nucleocapsid protein TR-FRET assay amenable to high-throughput screening"

Supplementary information

Supplementary Figure 1 - **Determination of NP concentration in viral samples**

Supplementary Figure 2 - **NP HTRF assay is able to detect recombinant SARS-CoV-2 NP**

Supplementary Figure 3 - **NP HTRF assay is able to detect SARS-CoV-2 NP in TCS and cell lysates**

Supplementary Figure 4 - **NP HTRF assay is able to detect transiently transfected SARS-CoV-2 NP**

Supplementary Figure 5 - **Sequence alignment of VoC Beta, Gamma, and Epsilon compared with SARS-CoV-2 USA-WA1/2020**

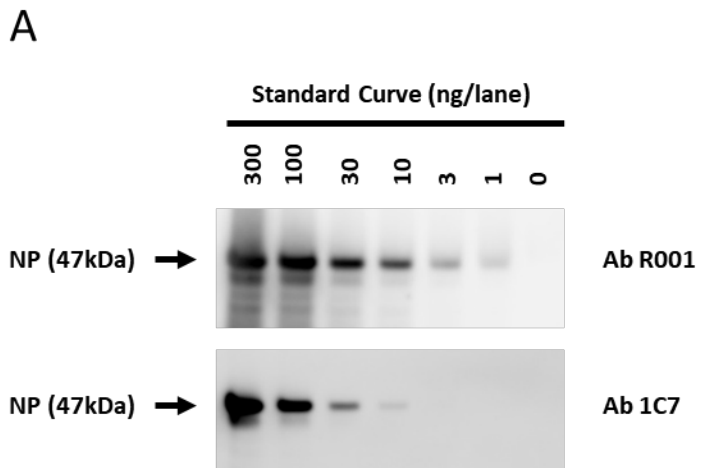


**Supplementary Figure 1. Determination of NP concentration in viral samples.** Western blot of recombinant SARS-CoV-2 NP using donor antibody R001 and acceptor antibody 1C7 at 1:1000 dilution.

A

Media only 1h

| | 1h incubation | | | |
|---|---|---|---|---|
| Concentration (ng/mL) | 5 nM D: 10 nM A | 1 nM D: 2 nM A | 10 nM D: 10 nM A | 2 nM D: 2 nM A |
| 1500 | 12.6 | 43.5 | 23.5 | 55.0 |
| 750 | 8.7 | 38.7 | 15.2 | 54.8 |
| 375 | 5.2 | 23.3 | 8.7 | 37.1 |
| 187.5 | 3.2 | 12.8 | 4.9 | 21.8 |
| 93.8 | 2.1 | 7.1 | 3.1 | 10.9 |
| 46.9 | 1.6 | 4.3 | 2.0 | 6.4 |
| 23.4 | 1.4 | 2.6 | 1.6 | 3.9 |
| 11.7 | 1.2 | 1.9 | 1.3 | 2.5 |
| 5.9 | 1.1 | 1.5 | 1.1 | 1.7 |
| 2.9 | 1.0 | 1.3 | 1.1 | 1.4 |
| 1.5 | 1.0 | 1.1 | 1.0 | 1.2 |
| 0 | 1.0 | 1.0 | 1.0 | 1.0 |

B

Media only O/N

| | O/N 4°C | | | |
|---|---|---|---|---|
| Concentration (ng/mL) | 5 nM D: 10 nM A | 1 nM D: 2 nM A | 10 nM D: 10 nM A | 2 nM D: 2 nM A |
| 1500 | 9.2 | 35.1 | 17.7 | 45.0 |
| 750 | 6.6 | 37.7 | 11.5 | 51.1 |
| 375 | 4.2 | 26.0 | 6.4 | 42.2 |
| 187.5 | 2.7 | 12.8 | 3.8 | 24.6 |
| 93.8 | 1.9 | 6.5 | 2.5 | 11.4 |
| 46.9 | 1.5 | 3.8 | 1.7 | 6.1 |
| 23.4 | 1.3 | 2.4 | 1.4 | 3.6 |
| 11.7 | 1.1 | 1.7 | 1.2 | 2.4 |
| 5.9 | 1.1 | 1.4 | 1.0 | 1.8 |
| 2.9 | 1.0 | 1.2 | 1.0 | 1.4 |
| 1.5 | 1.0 | 1.1 | 1.0 | 1.2 |
| 0 | 1.0 | 1.0 | 1.0 | 1.0 |

C

Media + cells 1h

| | 1h incubation | | | |
|---|---|---|---|---|
| Concentration (ng/mL) | 5 nM D: 10 nM A | 1 nM D: 2 nM A | 10 nM D: 10 nM A | 2 nM D: 2 nM A |
| 1500 | 15.0 | 39.9 | 23.8 | 45.2 |
| 750 | 11.3 | 36.5 | 19.0 | 50.8 |
| 375 | 7.2 | 29.5 | 12.4 | 39.6 |
| 187.5 | 4.6 | 18.5 | 7.1 | 27.6 |
| 93.8 | 2.9 | 11.1 | 4.3 | 15.2 |
| 46.9 | 2.0 | 6.0 | 2.7 | 10.1 |
| 23.4 | 1.5 | 3.8 | 1.9 | 5.1 |
| 11.7 | 1.2 | 2.5 | 1.5 | 3.8 |
| 5.9 | 1.1 | 1.9 | 1.2 | 2.5 |
| 2.9 | 1.0 | 1.4 | 1.1 | 1.7 |
| 1.5 | 1.0 | 1.2 | 1.0 | 1.4 |
| 0 | 1.0 | 1.0 | 1.0 | 1.0 |

D

Media + cells O/N

| | O/N 4°C | | | |
|---|---|---|---|---|
| Concentration (ng/mL) | 5 nM D: 10 nM A | 1 nM D: 2 nM A | 10 nM D: 10 nM A | 2 nM D: 2 nM A |
| 1500 | 12.0 | 34.7 | 22.4 | 36.8 |
| 750 | 9.1 | 42.8 | 16.6 | 48.0 |
| 375 | 5.6 | 37.2 | 9.1 | 49.3 |
| 187.5 | 3.7 | 20.8 | 5.4 | 35.1 |
| 93.8 | 2.5 | 10.6 | 3.4 | 17.7 |
| 46.9 | 1.7 | 5.7 | 2.2 | 9.5 |
| 23.4 | 1.4 | 3.3 | 1.6 | 5.3 |
| 11.7 | 1.2 | 2.3 | 1.3 | 3.3 |
| 5.9 | 1.0 | 1.6 | 1.1 | 2.3 |
| 2.9 | 1.0 | 1.3 | 1.0 | 1.6 |
| 1.5 | 0.9 | 1.2 | 1.0 | 1.4 |
| 0 | 1.0 | 1.0 | 1.0 | 1.0 |

**Supplementary Figure 2. NP HTRF assay is able to detect recombinant SARS-CoV-2 NP**. **(A)** S/B for Vero E6 NP in media with 1h incubation. **(B)** S/B for Vero E6 NP in media with O/N incubation. **(C)** S/B for Vero E6 NP in media and cells with 1h incubation. **(D)** S/B for Vero E6 NP in media and cells with O/N incubation.

A

| Dilution Factor | Mock TCS | 24h TCS | 48h TCS |
|---|---|---|---|
| no dilution | 0.8 | 5.4 | 9.8 |
| 15 | 1.1 | 2.0 | 8.1 |
| 45 | 1.0 | 1.4 | 3.7 |
| 135 | 1.1 | 1.1 | 2.0 |
| 405 | 1.0 | 1.0 | 1.4 |
| 1215 | 1.0 | 1.0 | 1.1 |
| 3645 | 1.0 | 1.0 | 1.1 |
| 10935 | 1.0 | 1.0 | 1.0 |
| 32805 | 1.0 | 1.0 | 1.0 |
| 98415 | 1.0 | 1.0 | 1.0 |
| 295245 | 1.0 | 0.9 | 1.0 |
| 885735 | 1.0 | 1.0 | 1.0 |

B

| Dilution Factor | Mock Lysate | 24h Lysate | 48h Lysate |
|---|---|---|---|
| 15 | 0.8 | 9.7 | 6.0 |
| 45 | 1.1 | 26.8 | 41.0 |
| 135 | 1.0 | 15.8 | 31.5 |
| 405 | 1.0 | 6.0 | 16.6 |
| 1215 | 1.0 | 2.6 | 6.2 |
| 3645 | 1.0 | 1.5 | 2.5 |
| 10935 | 1.0 | 1.2 | 1.4 |
| 32805 | 1.0 | 1.0 | 1.0 |
| 98415 | 1.0 | 1.0 | 1.0 |
| 295245 | 1.0 | 1.0 | 1.0 |
| 885735 | 1.0 | 0.9 | 0.9 |
| 2657205 | 1.0 | 1.0 | 1.0 |

**Supplementary Figure 3. NP HTRF assay is able to detect SARS-CoV-2 NP in TCS and cell lysates. (A)** S/B for Vero E6 TCS. **(B)** S/B for Vero E6 cell lysate.

A

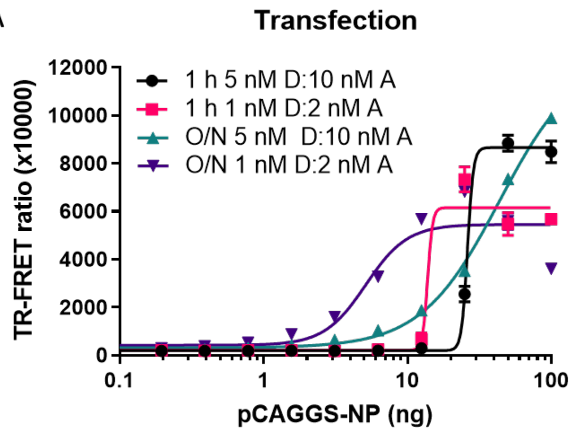


B

| | 1 h incubation | | O/N 4°C | |
|---|---|---|---|---|
| Concentration (ng) | 5 nM D: 10 nM A | 1 nM D: 2 nM A | 5 nM D: 10 nM A | 1 nM D: 2 nM A |
| 100.0 | 45.4 | 25.7 | 71.2 | 26.7 |
| 50.0 | 47.3 | 24.9 | 75.7 | 25.7 |
| 25.0 | 13.6 | 33.0 | 12.0 | 52.0 |
| 12.5 | 1.6 | 3.0 | 1.5 | 2.5 |
| 6.3 | 1.1 | 1.1 | 1.0 | 0.9 |
| 3.1 | 1.0 | 0.8 | 1.0 | 0.9 |
| 1.6 | 1.0 | 0.9 | 1.0 | 0.9 |
| 0.8 | 1.0 | 0.9 | 1.0 | 0.9 |
| 0.4 | 1.0 | 0.9 | 1.0 | 0.9 |
| 0.2 | 1.0 | 0.9 | 1.0 | 0.9 |
| 0.1 | 1.0 | 1.0 | 1.0 | 0.9 |
| 0.0 | 1.0 | 1.0 | 1.0 | 1.0 |

C

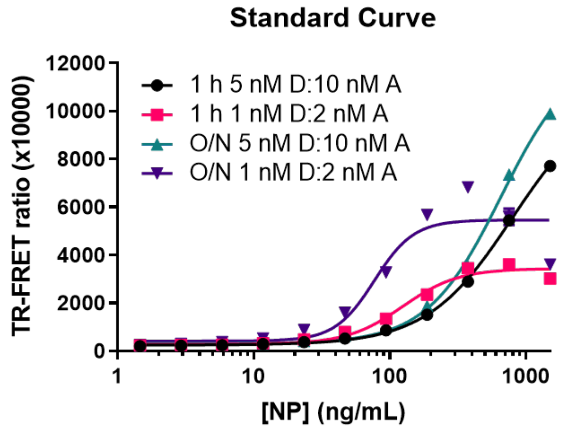


D

| | 1 h incubation | | O/N 4°C | |
|---|---|---|---|---|
| Concentration (ng/mL) | 5 nM D: 10 nM A | 1 nM D: 2 nM A | 5 nM D: 10 nM A | 1 nM D: 2 nM A |
| 1500 | 40.5 | 16.4 | 51.5 | 14.5 |
| 750 | 28.6 | 19.5 | 38.3 | 22.4 |
| 375 | 15.2 | 18.6 | 18.3 | 27.4 |
| 187.5 | 7.9 | 12.7 | 9.8 | 22.7 |
| 93.8 | 4.5 | 7.2 | 5.4 | 13.1 |
| 46.9 | 2.8 | 4.3 | 3.3 | 6.4 |
| 23.4 | 2.0 | 2.6 | 2.2 | 3.5 |
| 11.7 | 1.5 | 1.8 | 1.6 | 2.1 |
| 5.9 | 1.3 | 1.4 | 1.3 | 1.5 |
| 2.9 | 1.2 | 1.3 | 1.2 | 1.2 |
| 1.5 | 1.1 | 1.3 | 1.2 | 1.0 |
| 0 | 1.0 | 1.0 | 1.0 | 1.0 |

**Supplementary Figure 4. NP HTRF assay is able to detect transiently transfected SARS-CoV-2 NP. (A)** TR-FRET ratio from Vero E6 cells transfected with a pCAGGS plasmid encoding SARS-CoV-2 USA-WA1/2020 NP for 24h starting at 100 ng serially diluted 1:2. **(B)** S/B values for A. **(C)** TR-FRET ratio from Vero E6 cells treated with SARS-CoV-2 NP starting at 1500 ng/mL and serially diluted 1:2. **(D)** S/B values for C. Cells were incubated with HTRF reagents for 1h at RT or O/N at 4°C. N=3 wells in a half-area 96-well plate. Error bars indicate S.D.

```
                              P80R
                               ↓
USA/WA-1   61   KEDLKFPRGQGVPINTNSSPDDQIGYYRRATRRIRGGDGKMKDLSPRWYFYYLGTGPEAG 120
Beta       61   KEDLKFPRGQGVPINTNSSPDDQIGYYRRATRRIRGGDGKMKDLSPRWYFYYLGTGPEAG 120
Gamma      61   KEDLKFPRGQGVPINTNSSRDDQIGYYRRATRRIRGGDGKMKDLSPRWYFYYLGTGPEAG 120
Epsilon    61   KEDLKFPRGQGVPINTNSSPDDQIGYYRRATRRIRGGDGKMKDLSPRWYFYYLGTGPEAG 120

                          R203K G204R T205I                    M234I
                                 ↓                               ↓
USA/WA-1   181  QASSRSSSRSRNSSRNSTPGSSRGTSPARMAGNGGDAALALLLLDRLNQLESKMSGKGQQ 240
Beta       181  QASSRSSSRSRNSSRNSTPGSSRGISPARMAGNGGDAALALLLLDRLNQLESKMSGKGQQ 240
Gamma      181  QASSRSSSRSRNSSRNSTPGSSKRTSPARMAGNGGDAALALLLLDRLNQLESKMSGKGQQ 240
Epsilon    181  QASSRSSSRSRNSSRNSTPGSSKGISPARMAGNGGDAALALLLLDRLNQLESK I SGKGQQ 240
```

**Supplementary Figure 5. Sequence alignment of VoC Beta, Gamma, and Epsilon compared with SARS-CoV-2 USA-WA1/2020**. ClustalO alignment of amino acid sequences for NP regions that were different compared to the reference USA/WA-1 strain. Red, bolded, and underlined letters indicate mutations. Mutation P80R (Gamma), R203K (Gamma, Epsilon), G204R (Gamma), T205I (Beta, Epsilon), and M234I (Epsilon). Sequences were obtained from GISAID.
